# Expression levels of glycoprotein O (gO) vary between strains of human cytomegalovirus, influencing the assembly of gH/gL complexes and virion infectivity

**DOI:** 10.1101/299222

**Authors:** Le Zhang, Momei Zhou, Richard Stanton, Jeremy Kamil, Brent J. Ryckman

**Author notes:** <u>Corresponding author:</u> Dr. Brent J. Ryckman Division of Biological Sciences Interdisciplinary Science Building Rm. 215 The University of Montana Missoula, MT 59812.

## Abstract

Tropism of human cytomegalovirus (HCMV) is influenced by the envelope glycoprotein complexes gH/gL/gO and gH/gL/UL128-131. During virion assembly, gO and the UL128-131 proteins compete for binding to gH/gL in the ER. This assembly process clearly differs among strains since Merlin (ME) virions contain abundant gH/gL/UL128-131 and little gH/gL/gO, whereas TR contains much higher levels of total gH/gL, mostly in the form of gH/gL/gO, but much less gH/gL/UL128-131 than ME. Remaining questions include 1) what are the mechanisms behind these assembly differences, and 2) do differences reflect *in vitro* culture adaptations or natural genetic variations? Since the UL74(gO) ORF differs by 25% of amino acids between TR and ME, we analyzed recombinant viruses in which the UL74(gO) ORF was swapped. TR virions were >40-fold more infectious than ME. Transcriptional repression of UL128-131 enhanced infectivity of ME to the level of TR, despite still far lower levels of gH/gL/gO. Swapping the UL74(gO) ORF had no effect on either TR or ME. A quantitative immunoprecipitation approach revealed that gH/gL expression was within 4-fold between TR and ME, but gO expression was 20-fold less by ME, and suggested differences in mRNA transcription, translation or rapid ER-associated degradation of gO. Trans-complementation of gO expression during ME replication gave 6-fold enhancement of infectivity beyond the 40-fold effect of UL128-131 repression alone. Overall, strain variations in assembly of gH/gL complexes result from differences in expression of gO and UL128-131, and selective advantages for reduced UL128-131 expression during fibroblast propagation are much stronger than for higher gO expression.

**IMPORTANCE:** Specific genetic differences between independently isolated HCMV strains may result from purifying selection on *de novo* mutations arising during propagation in culture, or random sampling among the diversity of genotypes present in clinical specimens. Results presented indicate that while reduced UL128-131 expression may confer a powerful selective advantage during cell-free propagation of HCMV in fibroblast cultures, selective pressures for increased gO expression are much weaker. Thus, variation in gO expression among independent strains may represent natural genotype variability present *in vivo*. This may have important implications for virus-host interactions such as immune recognition, and underscores the value of studying molecular mechanisms of replication using multiple HCMV strains.

## INTRODUCTION

Human cytomegalovirus (HCMV) is widely spread throughout the world, found in approximately 60% of adults in developed countries and 100% in developing countries (reviewed in (1–3) (4)). Immunocompromised individuals such as those infected with HIV patients, or transplant recipients under antirejection treatments can suffer HCMV related pathologies including gastroenteritis, encephalitis, retinitis, and vasculopathies, which can accelerate allograft rejection. HCMV infection can also be acquired in utero and this is a significant cause of congenital neurological impairments and sensorineural hearing loss. The transmission of HCMV is mainly through body liquid, such as urine and saliva (5). Once infection is established, HCMV can spread throughout the body, infecting many of the major somatic cell types including, fibroblasts smooth muscle cells, epithelial and endothelial cells, neurons, and leukocytes such as monocytes-macrophages, and dendritic cells (6–9). HCMV does not replicate efficiently in transformed cells (10, 11), thus most studies of the mechanisms governing HCMV tropism have involved dermal fibroblasts, and retinal pigment epithelial cells and umbilical cord endothelial cells, all of which can be easily cultured as normal, non-transformed cells.

Much focus has been on the gH/gL complexes, which as for other herpesviruses, likely engage cell receptors and promote infection by contributing to the gB-mediated membrane fusion event or through activating cell signaling pathways (reviewed in (12, 13) (14)). During virus assembly, the HCMV UL128-131 proteins and gO compete for binding to gH/gL to form the pentameric complex gH/gL/UL128-131, or the trimeric complex gH/gL/gO. Structural studies involving purified soluble complexes showed that gO and UL128 can each make a disulfide bond with cystine 144 of gL, and this was suggested to be the basis of the competitive assembly of the complexes (15). However, Stegmann 2017 demonstrated that a mutant gO lacking the cysteine implicated in the disulfide bond with gL formed an intact, and functional gH/gL/gO (16). This suggests that gO can engage in extensive non-covalent interactions with gH/gL. The gH/gL/UL128-131 complex is dispensable for infection of cultured fibroblasts and neuronal cells, but required for infection of epithelial endothelial cells and monocyte-macrophages (17) (18) (19) (20) (21). In contrast, gH/gL/gO is critical for infection of all cell types (22) (23) (24) (25). Both complexes likely interact with cell receptors. gH/gL/gO can bind platelet-derived growth factor receptor-alpha (PDGFRα□ through the gO subunit, and this interaction is critical for infection of fibroblasts (26–28). Epithelial and endothelial cells do not express PDGFRα, but blocking of gH/gL/gO either with neutralizing antibodies or with soluble PDGFRα can inhibit infection of these cells, suggesting the existence of other gH/gL/gO receptors (26, 27). Receptors for gH/gL/UL128-131 might include epidermal growth factor receptor (EGFR; also known as ErbB1), and β1 or β3 integrins, and these interactions may induce signaling cascades, critical for infection of selected cell types such as epithelial and endothelial cells and monocyte-macrophages (26, 29).

We recently reported that the amounts of gH/gL/gO, and gH/gL/UL128-131 in the virion envelope can differ dramatically among commonly studied strains of HCMV, and that this can affect the infectivity of the virions (25, 30). The salient results of these studies were; 1) ME virions contained gH/gL mostly in the form of gH/gL/UL128-131, whereas TR and TB virions had mostly gH/gL/gO, 2) in terms of “total gH/gL”, the amount of gH/gL/gO in TR and TB virions was more than the gH/gL/UL128-131 in ME virions, 3) the infectivity of all three strains on both fibroblasts and epithelial cells correlated with the amount of gH/gL/gO, and 4) when the expression of UL128-131 was suppressed in ME, virions contained dramatically less gH/gL/UL128-131, but only slightly more gH/gL/gO. This latter point was especially curious since the model that gO and the UL128-131 proteins compete for binding to gH/gL would predict that the fraction of gH/gL normally bound by UL128-131 would, in their absence, be instead bound by gO. This discrepancy could be explained by differences in the stoichiometric expression of gH/gL, gO, and UL128-131 between strains. An alternative hypothesis was suggested by the fact that there are at least eight alleles of the UL74 gene that encodes gO (31). Isoforms of gO can vary between 10-30% of amino acids, and this could affect competition with UL128-131 for binding to gH/gL. Both of these non-mutually exclusive hypotheses were addressed in the experiments reported here.

## RESULTS

### Strains of HCMV display different patterns of glycoprotein expression and trafficking to virion assembly compartments

The dramatic differences in the composition of gH/gL complexes in TR and ME virions described in Zhou 2013/2015 (25, 30) suggested corresponding differences in glycoprotein expression, and/or trafficking of glycoproteins to virion assembly compartments (AC). To address these possibilities, cells were infected for 2 days (Fig 1A) or 5 days (Fig 1B) with TR or ME, and steady state amounts of viral proteins were compared by immunoblot. At 2 dpi, immediate-early (IE)-1/2 levels were similar for both TR and ME, consistent with an equal multiplicity of infection. At 5 dpi, the levels of the virion structural proteins MCP, gB, gH, and gL were also very similar between the two strains. In contrast, ME infected cells contained dramatically more of the UL128-131 proteins than TR. The UL148 protein was also included in these analyses because it was recently described as an ER chaperone protein that influences the ratio of gH/gL complexes (32). In TR-infected cells, an anti-UL148 antibody detected a prominent 35 kDa protein species, consistent with the previous description of the UL148 protein (32). This 35-kDa species was not detected in ME-infected cells. Instead, ME-infected cells contained two species that were less abundant, and of faster and slower electrophoretic motilities than the single UL148 species detected in TR-infected cells. The basis of the apparent size difference was not characterized, but could reflect differences in translational start/stop codon usage, splicing of the UL148 mRNA, or posttranslational modifications of the UL148 protein between strains. Overall, the pattern of expression of the UL128-131, and UL148 proteins correlated well with the previously described pentamer-rich nature of ME virions and the trimer-rich nature of TR virions (25, 30). Note that the expression of gO was not addressed in these analyses because the gO amino acid sequence differences between strains affects antibody recognition and precluded direct comparison (30).

**Figure 1.**
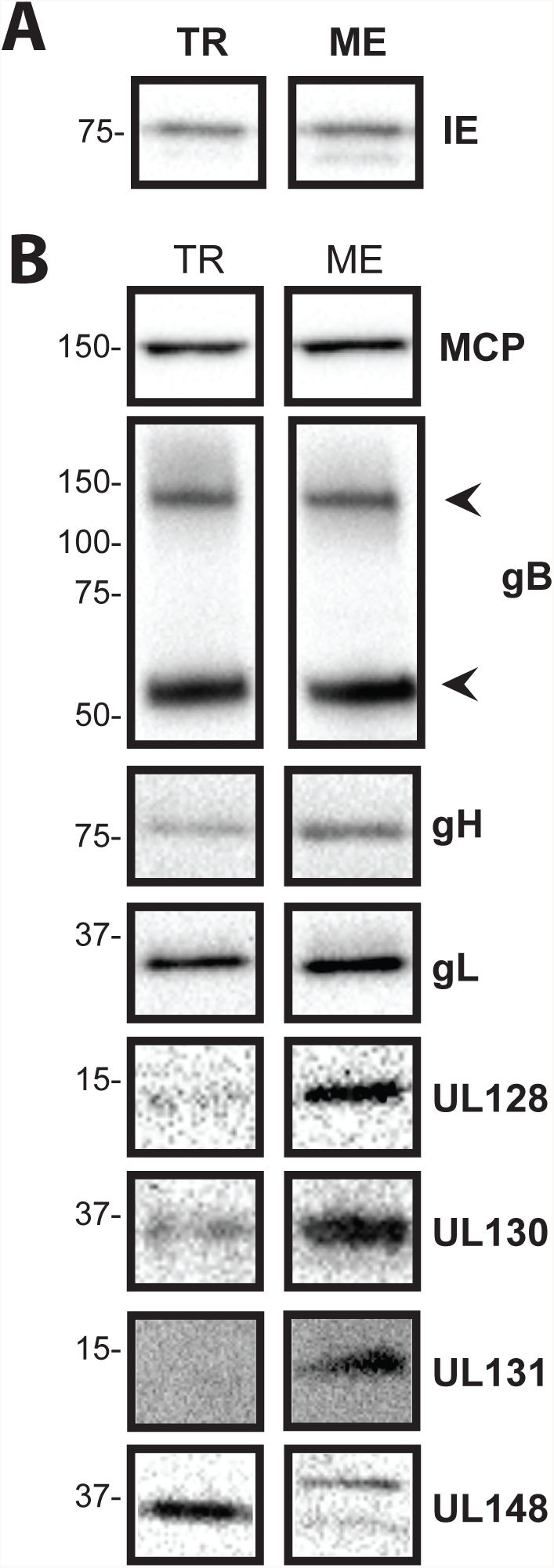
Comparison of protein expression between TR and ME. nHDF were infected with 1 PFU/cell of TR or ME. At day 2 (A) or day 5 (B) total cell extracts were separated by reducing SDS-PAGE and analyzed by immunoblot probing for immediate early (IE)-1/2, major capsid protein (MCP), gB, gH, gL, UL128, UL130, UL131, or UL148. Arrowheads indicate the positions of the cleaved 100kDa and 55 kDa fragments of gB.

Trafficking of gH/gL from the ER to TGN-derived assembly compartments was assessed by treating the 5 dpi infected-cell extracts with either endoglycosidase H (endo H) or PNGaseF, and then analyzing gH and gL by immunoblot (Fig 2). The majority of gH and gL in TR-infected cells was endo H resistant, consistent with efficient transport from the ER to *trans*-Golgi-derived ACs. In contrast, most of the gH and gL in ME-infected cells was sensitive to endo H digestion. In HFFFtet cells, which repress transcription from the UL128-131 locus (30) (33), there was even less endo H resistant gH and gL. This suggested that the bulk of gH/gL trafficked to ACs in ME-infected nHDF, which allow UL128-131 expression, represented gH/gL/UL128-131 and is consistent with the previous observations that, 1) the bulk of gH/gL in the ME virion is pentamer, and 2) the loss of gH/gL in the form of pentamer in ME-T virions due to the repression of the UL128-131 proteins is apparently not fully compensated by the formation of complexes with gO (25, 30).

**Figure 2.**
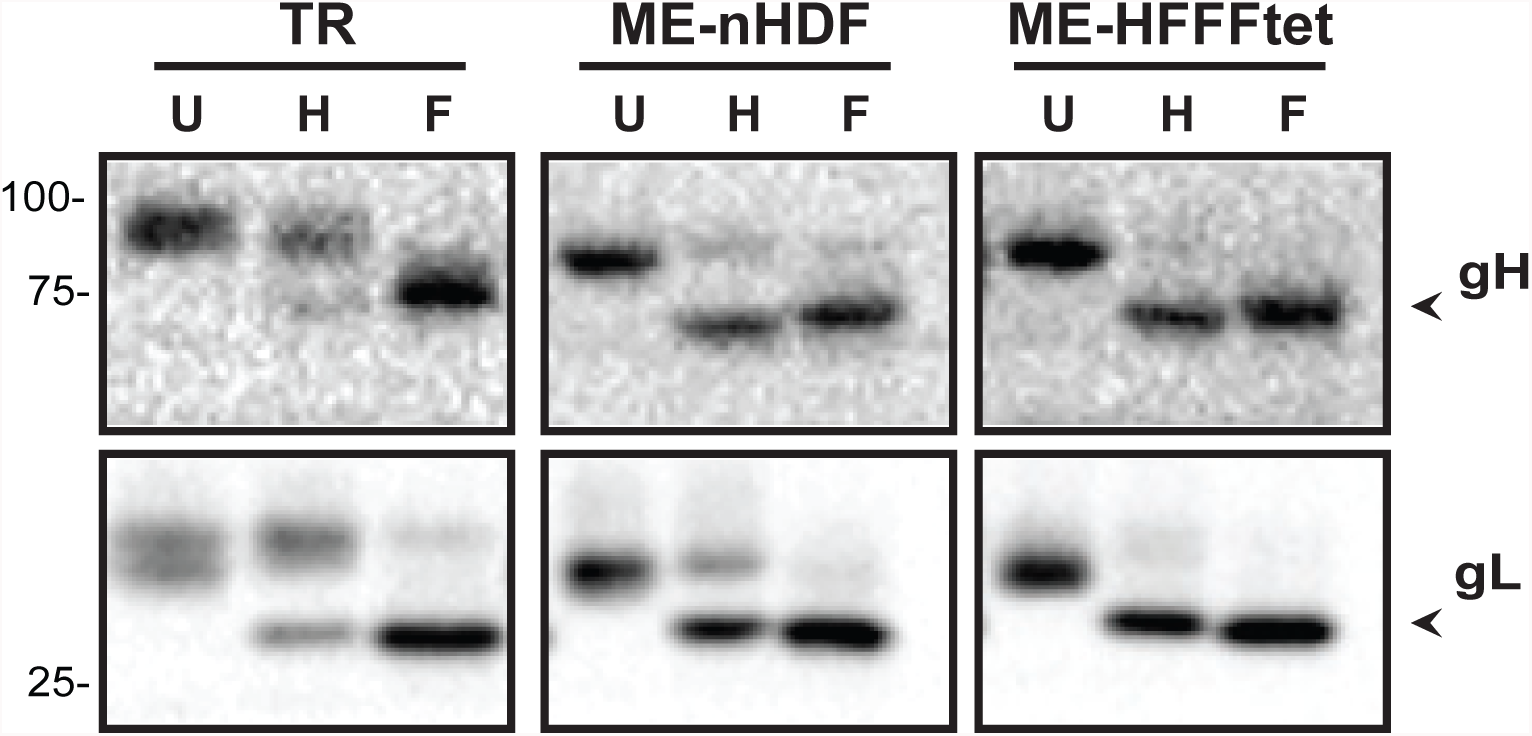
Analysis of ER-to-*trans*Golgi compartment trafficking of glycoproteins in TR or ME infected cells. Extracts of nHDF infected with TR or ME, or HFFFtet cells infected with ME were treated with endoglycosidase H (H), PNGaseF (F) or left untreated (U), and then separated by reducing SDS-PAGE and analyzed by immunoblot probing for gH or gL. Arrowheads indicate the position of the faster migrating, deglycosylated species.

### Differences in the amino acid sequence of gO between TR and ME do not affect the infectivity of cell free virus

The predicted amino acid sequence of gO differs by 25% between TR and ME. This sequence divergence precluded direct comparison of gO expression levels because antibodies do not cross-react (30). Furthermore, these sequence differences could potentially affect the ability of the distinct gO isoforms to compete with the UL128-131 proteins for binding to gH/gL (thus influencing the amounts of gH/gL complexes in the mature virion envelope), or the function(s) of gO during entry, such as binding PDGFRα or other receptors. To address these possibilities, BAC recombineering methods were used to replace the gO ORF (UL74) of TR with the analogous sequences from ME, and visa versa to generate recombinant viruses denoted TR_MEgO and ME_TRgO.

Zhou 2015 demonstrated a positive correlation between the infectivity of HCMV virions and the amounts of gH/gL/gO in the virion envelope (25). To assess the effects of gO sequences on infectivity, cell free virus stocks of parental wild type and heterologous gO recombinants were analyzed by qPCR to determine the number of virions, and infectivity was determined by plaque assay. No difference in particles/PFU was observed between TR and the corresponding recombinant, TR_MEgO (Fig 3A), or between ME and the corresponding recombinant ME_TRgO (Fig 3B). When the ME-based HCMV were grown in HFFFtet cells, which repress UL128-131 expression, the resultant virions, ME-T and ME-T_TRgO, were dramatically more infectious, as shown before (25) (33), but consistently there were no differences due to the isoform of gO expressed (Fig. 3B). In parallel analyses, the amounts of gH/gL complexes were analyzed by non-reducing immunoblot probing for gL to detect intact, disulfide linked gH/gL/gO, and disulfide-linked gH/gL/UL128 (note that UL130 and UL131 are not disulfide-linked to the intact pentamer complex and are thus separated by SDS-PAGE) (Fig 4). Consistent with our previous reports (25, 30), TR virions contained much greater amounts of total gH/gL, mostly in the form of gH/gL/gO, whereas ME virions contained less gH/gL, mostly as gH/gL/UL128-131. Repression of the UL128-131 proteins (ME-T) drastically reduced the amount of gH/gL/UL128-131, and increased the amount of gH/gL/gO. However, note that the amount of gH/gL/gO in ME-T virions was still less than the gH/gL/UL128-131 in ME virions, indicating that the repression of UL128-131 was not fully compensated by gO. In no case did expression of the heterologous gO isoform detectably influence the amounts of gH/gL complexes in HCMV virions. Together these results suggest that the amino acid sequence differences between TR and ME gO do not influence gH/gL complex assembly, or the function of gO in entry into fibroblasts.

**Figure 3.**
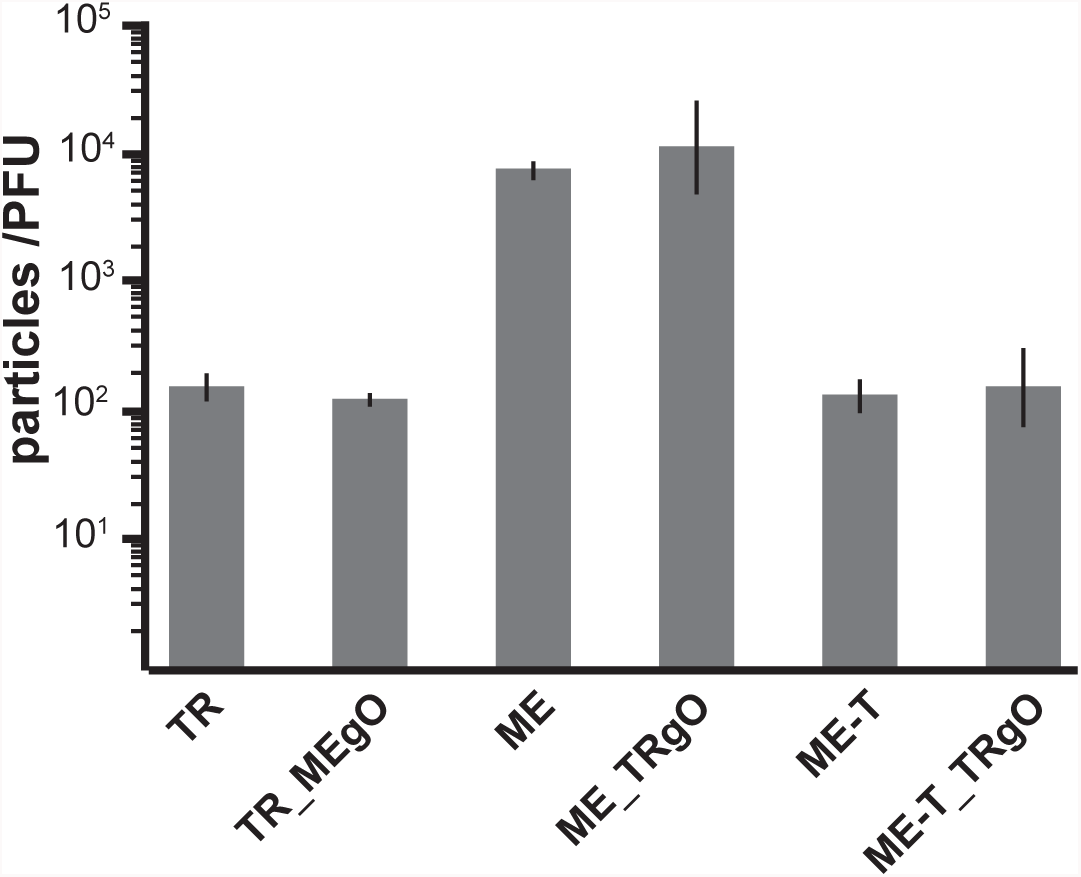
Specific infectivity of parental and TR-ME heterologous gO recombinants. Extracellular virions of TR, TR_MEgO, ME, ME_TRgO, ME-T, or ME-T_TRgO were analyzed by quantitative PCR for viral genomes and PFU were determined by plaque assay on nHDF. Average particle-PFU ratios from at least 4 independent experiments are plotted. Error bars represent standard deviation.

**Figure 4.**
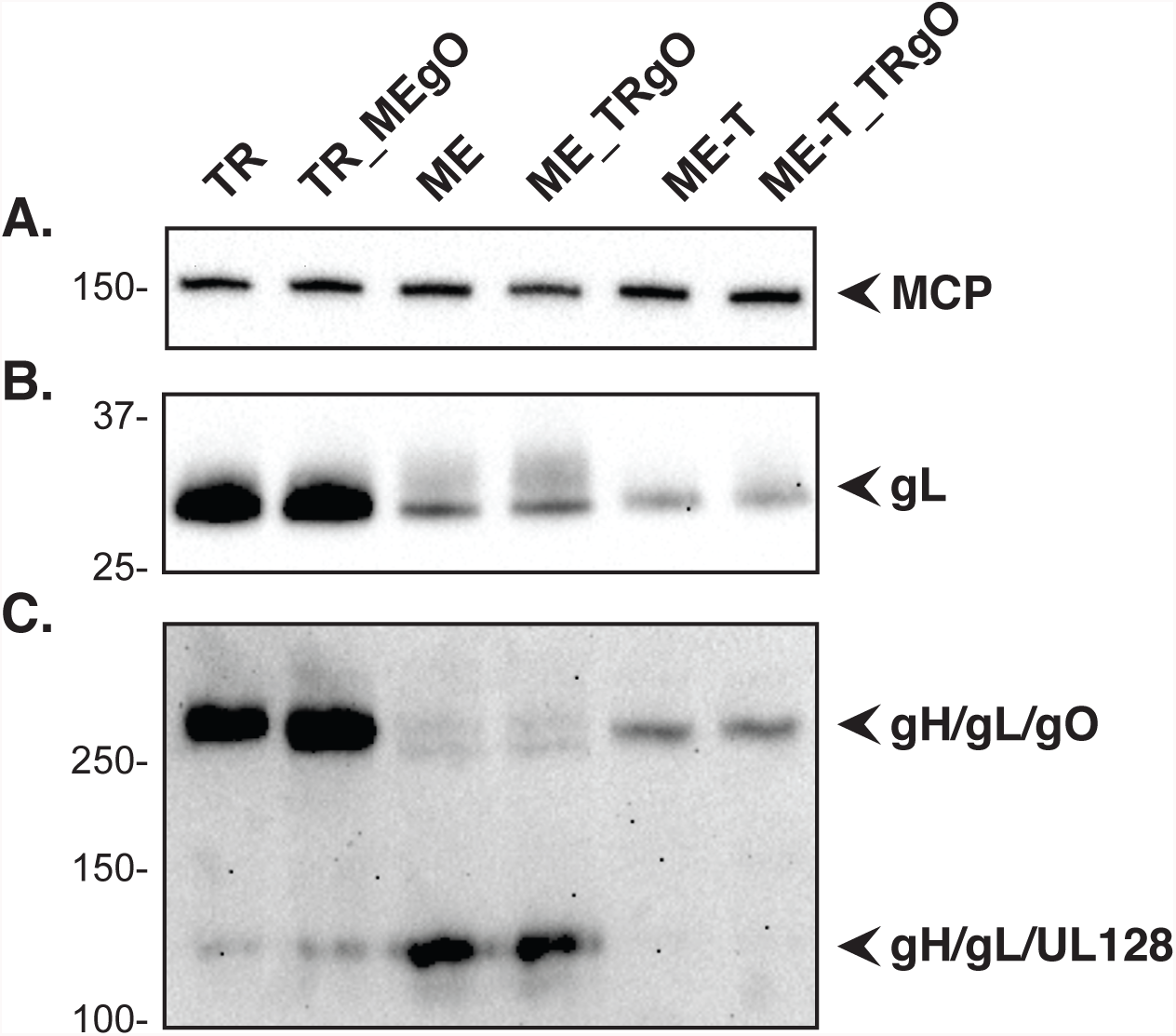
Immunoblot analysis of gH/gL complexes in parental and TR-ME heterologous gO recombinants. Extracellular virion extracts of TR, TR_MEgO, ME, ME_TRgO, ME-T, or ME-T_TRgO were separated by reducing (A and B) or non-reducing (C) SDS-PAGE and analyzed by immunoblot probing for major capsid protein (A) or gL (B and C)

### ME expresses less gO during replication than TR

The heterologous gO recombinants allowed comparison of gO expression level between TR and ME. In the first analyses cells infected with parental or the heterologous gO recombinants were analyzed by reducing immunoblot using TR and ME specific anti-gO antibodies (30) (Fig 5). TR-specific gO antibodies detected two bands in TR-infected cells, a prominent species migrating just above the 100kDa marker, and a minor, more diffuse species migrating at approximately 130-140kDa. The ME-specific antibodies detected similarly migrating bands in TR_MEgO infected cells, however their relative abundance appeared more equal. No similar bands were detected in cells infected with ME or ME_TRgO analyzed with either gO antiserum. The failure to detect either isoform of gO in cells infected with ME-based HCMV suggested that protein expression from the UL74 locus of ME was lower than in TR.

**Figure 5.**
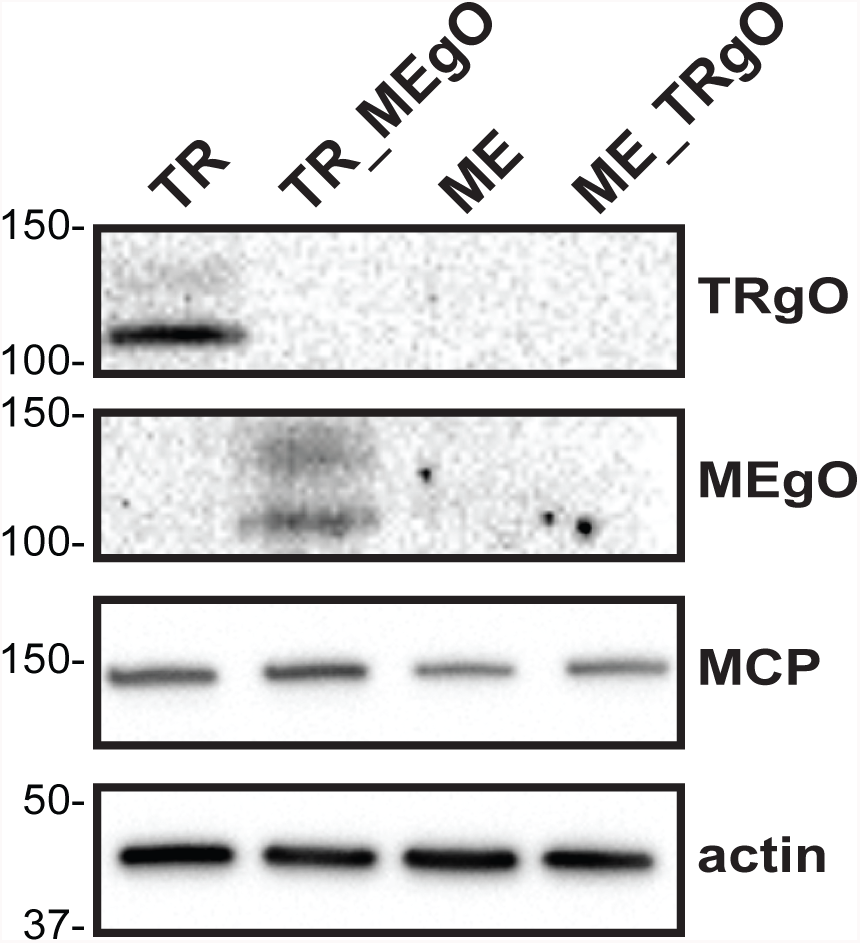
Immunoblot analysis of gO expression in cells infected with parental and TR-ME heterologous gO recombinants. nHDF were infected with 1 PFU/cell of TR, TR_MEgO, ME, or ME_TRgO. At day 5, total cell extracts of infected cells were separated by reducing SDS-PAGE and analyzed by immunoblot probing for TRgO, MEgO, major capsid protein (MCP), or actin.

To directly compare differences in glycoprotein expression between TR and ME, infected cells were labeled with [35]S-methionine/cysteine for 15 min, then analyzed by immunoprecipitation with anti-peptide antibodies specific for gH, gL, or gO, followed by SDS-PAGE and band density analysis (Fig 6, Tables 1 and 2). Two approaches were taken to allow for direct quantitative comparisons of labeled-proteins between extracts. First, cell extracts were denatured and reduced with SDS/DDT prior to immunoprecipitation to allow maximum epitope access by the anti-peptide antibodies. Second, for each analysis, multiple immunoprecipitation reactions were performed in parallel with increasing amounts of protein extract input to insure that antibodies were not limiting. In these experiments, expression of gH was nearly identical between TR and ME, gL expression was approximately 4-fold higher for TR than for ME, but gO expression was strikingly 27-fold higher for TR than for ME (Fig. 6A, Table 1). To address the possibility that the MEgO-specific antibodies were simply less efficient at capturing MEgO from ME extracts, similar experiments were performed with the TR-ME heterologous gO recombinants (Fig 6B, Table 2). Again, gH and gL were similar between TR_MEgO and ME_TRgO, but gO levels were approximately 20-fold lower higher for the TR-based virus. To address the hypothesis that differences in gO expression between TR and ME reflect differences in protein turnover, the [35]S-methionine/cysteine label was chased for up to 6 hours (Fig 7). The pattern of gH detection over the chase time was very similar in both TR and ME samples. In both cases, labeled gH dropped to 60% after 3 hours and to 30-40% after 6 hours. The pattern of gO detection for both TR and ME was comparable to that of gH. Together, these results confirmed that ME-infected cells express less gO than TR-infected cells, and suggested differences in early steps of expression such as mRNA transcription, translation or rapid ER-associated degradation, which can degrade proteins in the timescale of minutes (34).

**Table 1.** Quantitative comparison of glycoprotein expression in TR and ME infected cells.

| IP Antibody<sup>a</sup> | Extract<sup>b</sup> (mL) | Strain |  |  |  | Fold<sup>f</sup> | Mean Fold<sup>g</sup> |
| --- | --- | --- | --- | --- | --- | --- | --- |
|  |  | TR |  | ME |  |  |  |
|  |  | Density<sup>c</sup> | Adjusted Density<sup>e</sup> | Density | Adjusted Density |  |  |
| anti-gH | 0.04 | n.d.<sup>d</sup> | n.d. | n.d. | n.d. |  |  |
|  | 0.13 | 136.7 | 4.6 | 106.6 | 3.4 | 1.3 |  |
|  | 0.40 | 476.9 | 15.9 | 337.8 | 10.9 | 1.5 |  |
|  | 1.20 | 1200.7 | 40.0 | 872.9 | 28.2 | 1.4 | 1.4(+/-0.1) |
| anti-gL | 0.04 | 143.8 | 11.1 | n.d. | n.d. | n.d. |  |
|  | 0.13 | 679.1 | 52.2 | 127.2 | 9.8 | 5.3 |  |
|  | 0.40 | 1627.3 | 125.2 | 509.3 | 39.2 | 3.2 |  |
|  | 1.20 | 6071.3 | 467.0 | 1809.9 | 139.2 | 3.4 | 4.0 (+/-1.2) |
| anti-gO | 0.04 | 267.0 | 12.1 | n.d. | n.d. | n.d. |  |
|  | 0.13 | 1008.6 | 45.8 | n.d. | n.d. | n.d. |  |
|  | 0.40 | 3805.4 | 173.0 | 120.8 | 5.0 | 34.4 |  |
|  | 1.20 | 9251.8 | 420.5 | 478.9 | 20.0 | 21.1 | 27.2 (+/-9.4) |

a. 7uL of rabbit anti-peptide serum per immunoprecipitation reaction
b. Preparation of radiolabeled cell extracts described in legend to Figure 6, and in materials and methods.
c. Densities of bands shown in Figure 6A as determined using Image J v. 1.48
d. Band density not detected.
e. Density divided by the predicted number of methionine and cysteine residues: TRgH(30), MEgH(31), TRgL(13), MEgL(13), TRgO(22), MEgO(24)
f. Adjusted density TR divided by adjusted density ME.
g. Average fold difference between TR and ME +/- standard deviation.

**Table 2.**
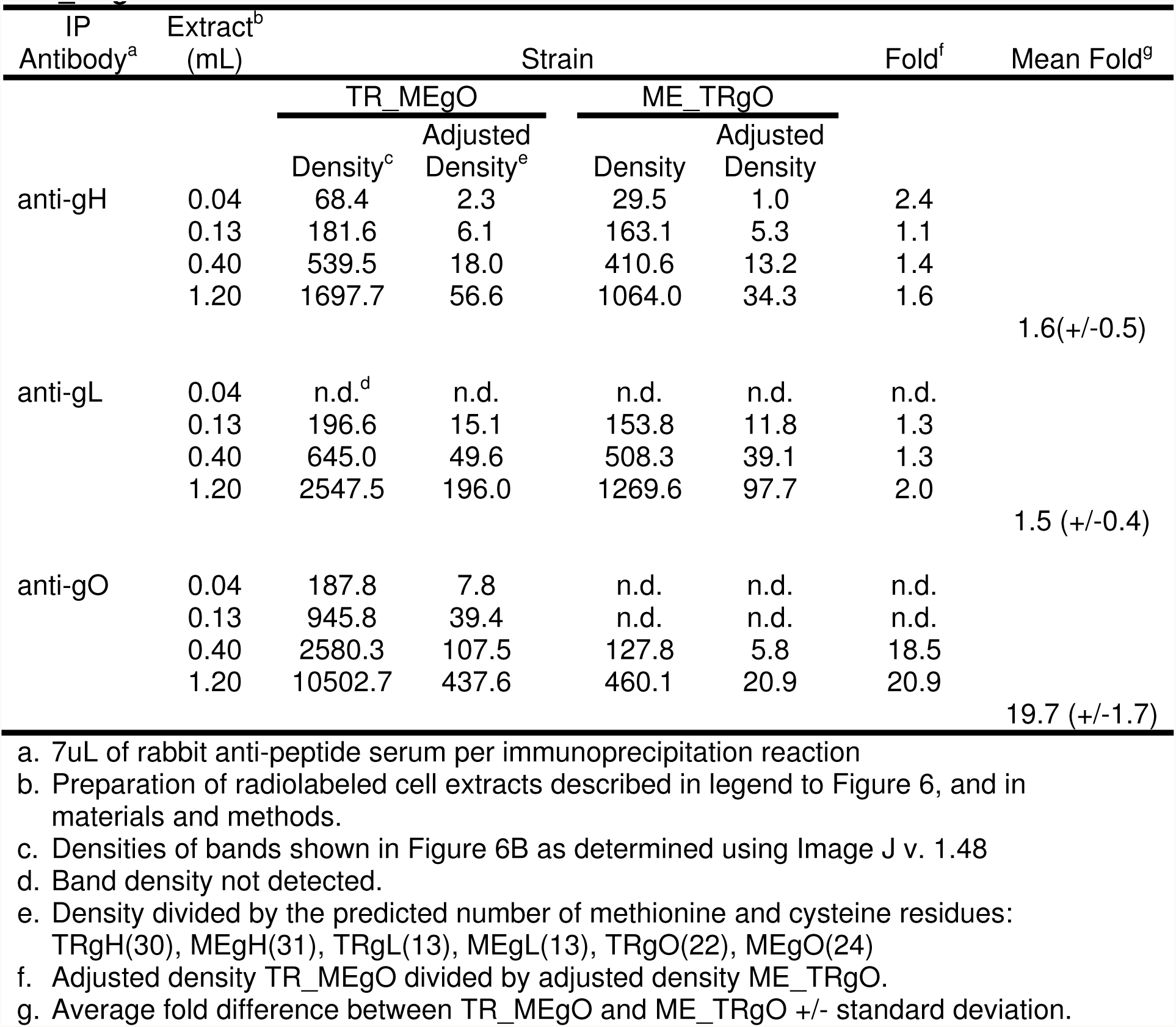
Quantitative comparison of glycoprotein expression in TR_MEgO and ME_TRgO infected cells.

**Figure 6.**
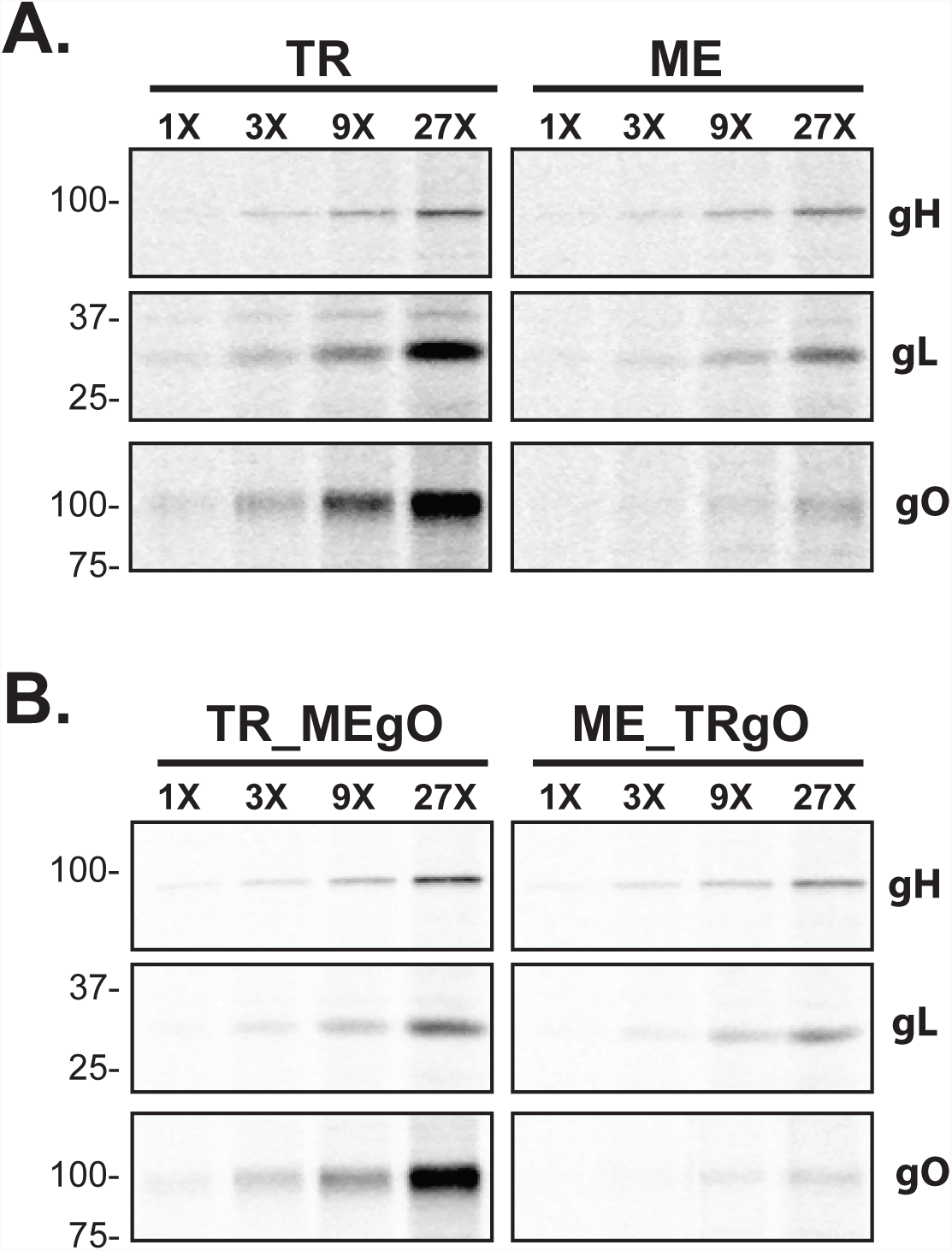
Quantitative comparison of glycoprotein expression in TR and ME infected cells. nHDF were infected with 1 PFU/cell of TR or ME (A), or TR_MEgO or ME_TRgO (B). At 5 dpi, infected cells were metabolically labeled with [35]-S cysteine/methionine for 15 min and membrane proteins were extracted in 1% Triton X-100. All samples were adjusted to 2%SDS/30mM DTT, heated to 75° C for 10 min, cooled to room temperature and then diluted 35-fold. Parallel immunoprecipitations were performed in which equal amounts of anti-gH, gL, or gO (TR or ME specific) antibodies were reacted with 3-fold increasing amounts of protein extract as input, and precipitated proteins were analyzed by SDS-PAGE.

**Figure 7.**
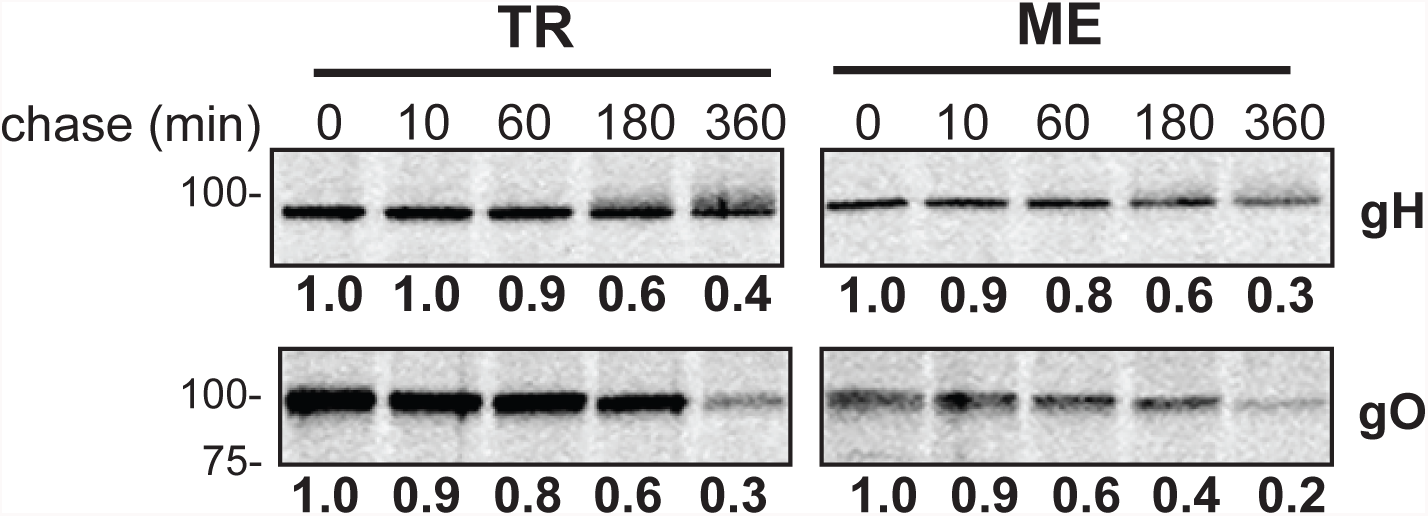
Analysis of glycoprotein turnover in TR and ME infected cells. nHDF were infected with 1 PFU/cell of TR or ME. At 5 dpi, infected cells were metabolically labeled with [35]-S cysteine/methionine for 15 minutes and then label was chased for 0, 10, 60, 180, or 360 minutes. Membrane proteins were extracted in 1% Triton X-100, adjusted to 2%SDS/30mM DTT, heated to 75° C for 10 min, cooled to room temperature and then diluted 35-fold. Immunoprecipitation was performed with anti-gH, gO (TR or ME specific) antibodies and precipitated proteins were analyzed by SDS-PAGE. Band densities were determined relative to the 0 minute chase time. Results shown are representative of 4 independent experiments.

### Overexpression of gO during ME replication increased gH/gL/gO assembly and virus infectivity

To directly test the hypothesis that the low abundance of gH/gL/gO in ME virions was due not simply to competition from the UL128-131 proteins, but also from low gO expression, Ad vectors were used to increase gO levels during ME replication. Ad vectors expressing GFP were used to control for potential effects of the Ad vectors themselves. Consistent with the above analyses, gO levels were below the limits of immunoblot detection in ME-infected nHDF or HFFF-tet cells, but gO was readily detected in cells superinfected with AdMEgO (Fig 8A). The overall expression of gL in ME infected cells was reduced by the presence of either Ad vector (Fig 8A). In the case of the control AdGFP, the lower intracellular gL correlated with reduced gH/gL/gO complexes in virions from HFFFtet cells (ME-T) (Fig 8B), and this in turn correlated with reduced infectivity (i.e., increased particle PFU ratio) (Fig 9). The “Ad effect” on virion gH/gL levels and infectivity was less apparent in HFF cells (ME), perhaps masked by the overall higher amounts of gH/gL and much lower infectivity of these virions (Fig 8B, and Fig. 9). Controlling for the “Ad effect”, AdMEgO expression in HFFFtet increased the amounts of gH/gL/gO in ME-T virions compared to the AdGFP, and this resulted in a 6-fold enhancement of infectivity, beyond the 40-fold enhanced infectivity resulting from repression of UL128-131 alone (Fig 8B, and Fig. 9). By contrast, AdMEgO expression had little effect on the virions from HFF cells.

**Figure 8.**
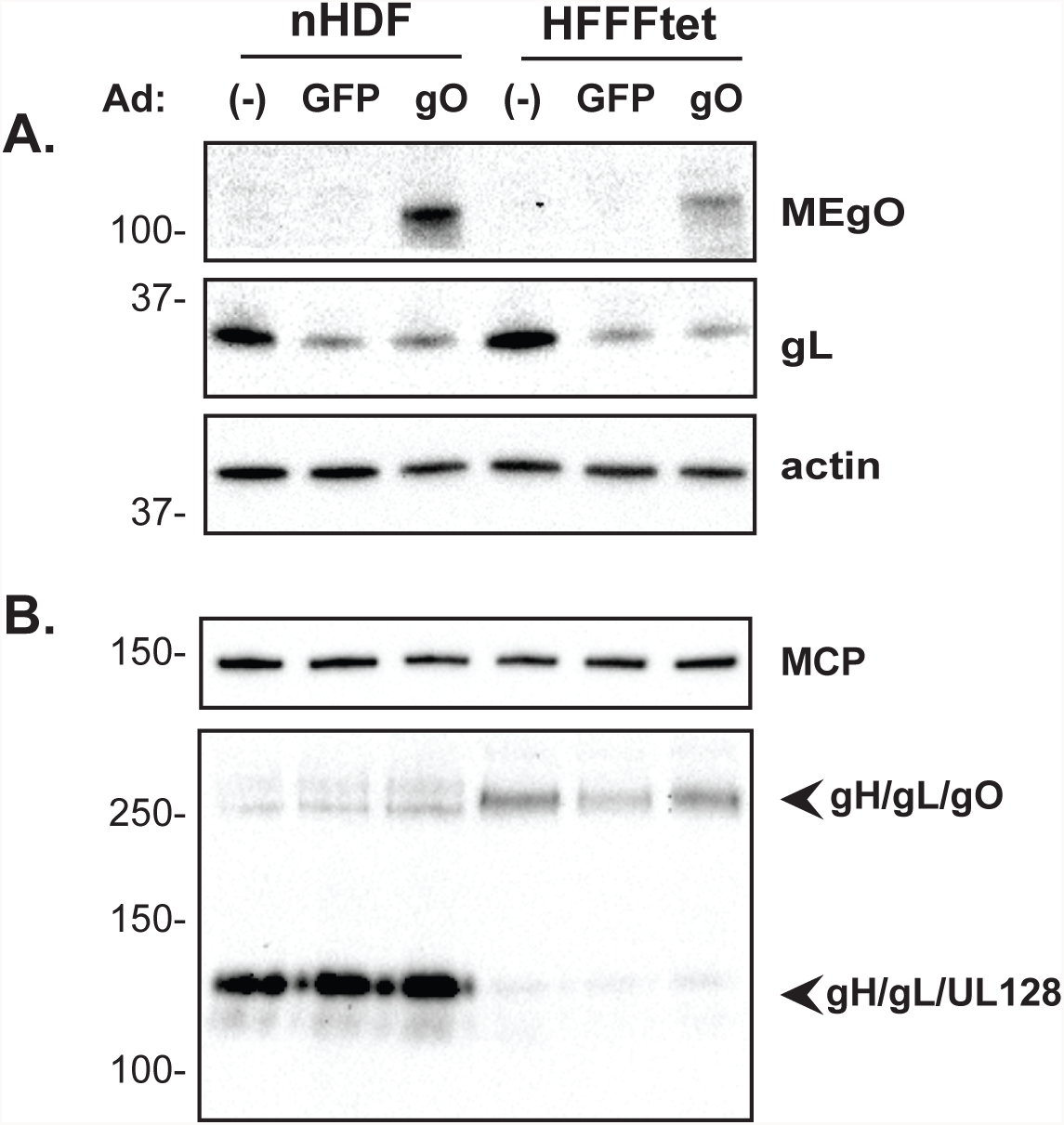
Ad vector overexpression of gO during ME replication. nHDF or HFFF-tet cells were infected with ME for 2 days, then superinfected with Ad vectors expressing either GFP or MEgO for an additional 4 days. Extracts of infected cells (A), or extracellular virions (B) were separated by reducing (A, and B; top) or non-reducing (B; bottom) SDS-PAGE and analyzed by immunoblot probing for MEgO, actin, major capsid protein (MCP), or gL as indicated to the right.

**Figure 9.**
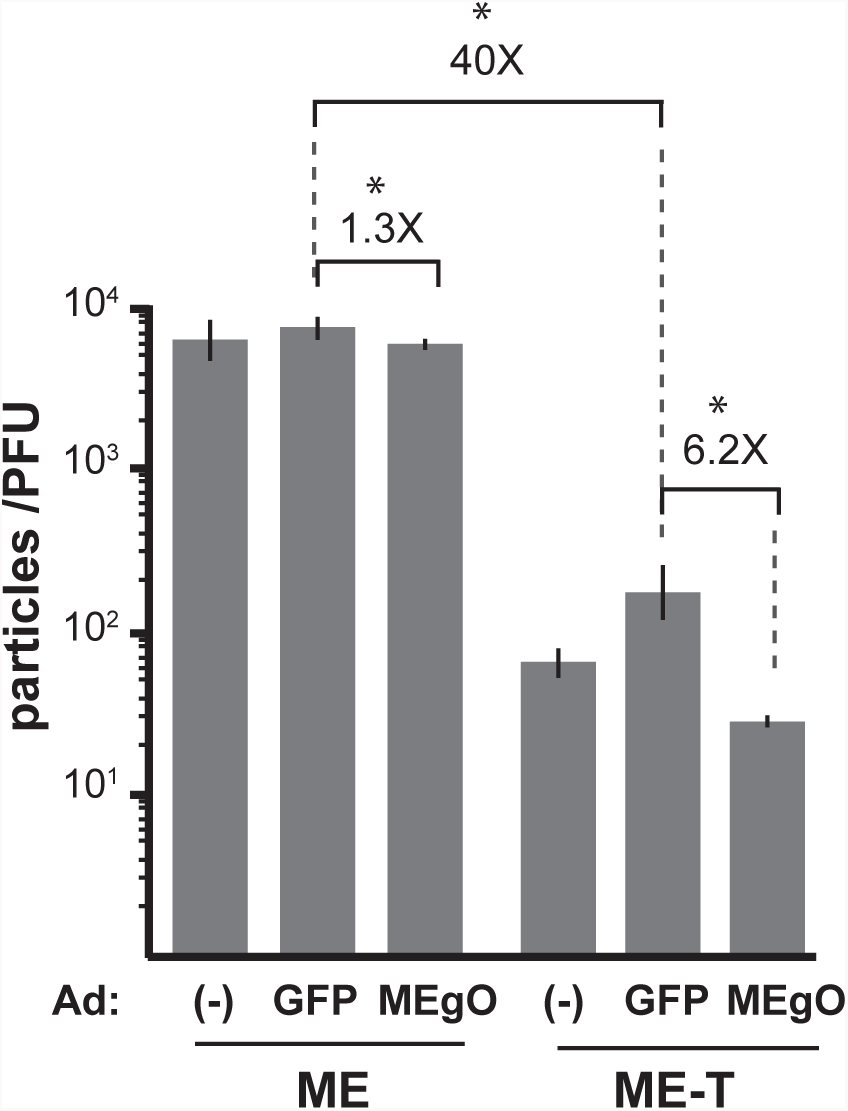
Specific infectivity of ME virions produced under conditions of gO overexpression. nHDF or HFFF-tet cells were infected with ME for 2 days, then superinfected with Ad vectors expressing either GFP or MEgO for an additional 4 days. Extracellular virions from nHDF (ME) or HFFFtet (ME-T) were analyzed by quantitative PCR for viral genomes, and PFUs determined by plaque assay on nHDF. Shown are average particle-PFU ratios of virions produced in 2 independent experiments, each analyzed in triplicate. Error bars represent the standard deviation. Asterisks (*) above fold differences indicate P < 0.03 (Student’s unpaired T-test (2-tailed)).

## DISCUSSION

Recent population genetic studies have demonstrated a greater degree of genetic diversity of HCMV in clinical specimens than had been previously appreciated (35, 36) (37). The cell type and propagation methods likely narrow the resultant genotypes by purifying selection (38, 39). During propagation in cultured fibroblasts, inactivating mutations in the UL128-131 ORFs are rapidly selected in a BAC clone of ME, and this selective pressure can be relieved by transcriptional repression of the UL131 promoter, which reduces expression of the pentameric gH/gL/UL128-131 (33) In contrast, the UL128-131 ORFs are more stable in BAC clones of strain TR, and TB (39, 40). The UL128-131 ORF of TB contains a single nucleotide polymorphism relative to ME that reduces splicing of the mRNA encoding the UL128 protein, which may help stabilize the UL128-131 ORFs through reduced expression of gH/gL/UL128-131 (40). However, TR is identical to ME at this nucleotide position, and a recombinant ME in which the UL128-131 locus was replaced with the UL128-131 sequences from TR was as sensitive to selective inactivation of the locus as was wild type ME (40). Together, these observations suggest that factors beyond expression level of the UL128-131 proteins can influence the selective pressures on the UL128-131 ORFs.

The results reported here demonstrated that TR and ME differ in stoichiometry of expression of gO and UL128-131, and this seems to be a major factor determining the abundance of gH/gL/gO and gH/gL/UL128-131 in the virion envelope, and the infectivity of cell free virion. Fibroblasts infected with TR or ME were found to be comparable in the steady state levels of gH/gL, but ME-infected cells contained more UL128-131 than TR infected cells. In ME-infected cells, most of the gH/gL was an ER-associated form, whereas TR-infected cells contained a large amount of Golgi-associated gH/gL. This correlated well with the previous observations that TR contained more total gH/gL than ME virions (25, 30). The amount of Golgi-associated gH/gL in ME–infected cells was reduced when expression of the UL128-131 proteins was repressed, consistent with the observation that most of the gH/gL in ME virions was in the form of gH/gL/UL128-131 (25, 30). Comparison of gO expression between strains was complicated because the amino acid sequence differences between genotypes affected antibody recognition (30). To circumvent this caveat, recombinant HCMV were engineered in which the UL74(gO) ORF of TR were replaced with the homologous sequences of ME, and *vice versa*. This approach allowed the analysis of expression of both gO isoforms in both genetic backgrounds eliminating the possibility that the results were due to differences in antibody-antigen affinities. Immunoblot and radiolabeling experiments clearly demonstrated that ME-infected cells contained less gO than TR-infected cells. Overexpression of gO during ME replication had no effect on levels of gH/gL/gO, or infectivity of the virions unless UL128-131 proteins were also transcriptionally repressed, and even then gH/gL/gO levels and infectivity were only modestly enhanced. Together these results underscore the competition between gO and UL128-131 for binding to gH/gL, and suggest other factors may influence the efficiency of gH/gL/gO assembly.

The molecular mechanisms underpinning the discrepancy between TR and ME in expression UL128-131, and gO remain unclear. As mentioned above, Murrell 2013 described a SNP in the TB UL128-131 locus that affected mRNA splicing, in part explaining the lower expression of these proteins in TB (40). However, this splicing effect does not explain the difference in UL128-131 expression between TR and ME since these strains are conserved at this nucleotide position. For gO, the radiolabeling analyses reported in Figures 6 and 7, suggest the differences are due to early events in UL74(gO) expression such as transcription, mRNA processing/stability, translation, or rapid ER-associated degradation occurring in the timescale of minutes (34). Attempts to analyze UL74(gO) mRNA levels between TR and ME by quantitative RT-PCR were complicated by the fact that HCMV genomes contain many overlapping RNAPII transcription units that vary between strains (41, 42). It is interesting that ME-infected cells contained less UL148 than TR-infected cells. UL148 was first described as an ER-resident chaperone protein that promotes the assembly of gH/gL/gO (32). The mechanism may well involve interactions between UL148 and the cellular ER-associated degradation pathway (C. Nguyen, M. Siddiquey, H.Zhang, J. Kamil; presented at 42^nd^ International Herpesvirus Workshop, 2017 Ghent Belgium).

The TR-ME heterologous gO recombinant viruses also allowed analysis of the effects of gO amino acid sequence differences on assembly of gH/gL complexes and the function of gO in entry. No differences between TR and TR_MEgO or between ME and ME_TRgO were observed in either the amounts of gH/gL complexes in virions, or cell free infectivity. These results argue against the notion that the amino acid sequence differences between gO genotypes affect interactions with gH/gL or the binding of the fibroblast entry receptor, PDGFRα. Interestingly Kalser 2017 showed that replacing the endogenous gO of TB with the gO from Towne did not alter replication in cultured fibroblasts, but did enhance replication in epithelial cell cultures. (43). Thus, it may be that gO sequence variation affects interactions with receptors other than PDGFRα that mediated infection of epithelial cells.

Laib-Sampaio 2016 reported that mutational disruption of UL74(gO) expression in ME had little effect on replication unless the UL128-131 locus was also disrupted (24). These authors suggested that spread of ME was mediated principally by gH/gL/UL128-131 in a cell-associated manner, but when UL128-131 was inactivated, spread could also occur in a cell-free manner mediated by gH/gL/gO. This is in stark contrast to the dramatic phenotype reported for a gO null TR mutant (22). Our finding that expression of gO by ME is low compared to TR may provide a partial explanation of these different gO null phenotypes.

It remains unclear whether the described difference in gO expression between TR and ME represents a *bona fide* variation that naturally exists between HCMV genotypes *in vivo*, or reflects differential selection on *de novo* mutations that occurred during the independent isolation of these strains from clinical specimens. It seems clear that serial propagation of ME in cultured fibroblasts selects for *de novo* mutations that reduce or abolish the robust expression of the UL128-131 proteins (33, 39). The selective pressure that fixes these mutations in the culture population may be explained by the specific infectivity analyses reported here (Figs. 3 and 9) and in Zhou 2015 (25). In both analyses the specific infectivity of TR was measured at approximately 100-200 particles/PFU, whereas ME was more 30-50-fold less infectious. Repression of the UL128-131 proteins enhanced the infectivity of ME (“ME-T”) to levels comparable to TR (approximately 100 particles/PFU). While the infectivity of ME-T and TR virions was comparable, ME-T virions still contained far less gH/gL/gO than TR (Fig 4, and ((25)). Ad vector overexpression of gO enhanced infectivity of ME only 6-fold beyond the enhancement due to UL128-131 repression alone (Fig. 8 and 9). Together, these observations would seem to suggest that *in vitro* selective pressures for reduced UL128-131 expression are much more pronounced than any for enhanced gO expression. Thus, it is possible that the difference in gO expression between HCMV TR and ME derives not from selection on *de novo* mutations occurring during propagation in culture, but from nonselective, random sampling of the multitude of different genotypes that likely preexist in clinical specimens (35, 36) (37). Distinguishing these possibilities will require clear identification of the genomic sequences that determine gO expression level.

## MATERIALS AND METHODS

### Cell lines

Primary neonatal human dermal fibroblasts (nHDF; Thermo Fisher Scientific), MRC-5 fibroblasts (American Type Culture Collection; CCL-171), and HFFFtet cells (which express the tetracycline (Tet) repressor protein provided by Richard Stanton (Cardiff University, Cardiff, United Kingdom) (33) were grown in Dulbecco’s modified Eagle’s medium (DMEM: Thermo Fisher Scientific) supplemented with 6% heat-inactivated fetal bovine serum (FBS; Rocky Mountain Biologicals, Inc., Missoula, Montana, USA.) and 6% bovine growth serum (BGS; Rocky Mountain Biologicals, Inc., Missoula, Montana, USA.).

### Human cytomegaloviruses

All HCMV were derived from bacterial artificial chromosome (BAC) clones. The BAC clone of TR was provided by Jay Nelson (Oregon Health and Sciences University, Portland, OR, USA) (44). The BAC clone of Merlin (ME) (pAL1393), which carries tetracycline operator sequences in the transcriptional promoter of UL130 and UL131 was provided by Richard Stanton (Cardiff University, Cardiff, United Kingdom) (33). Infectious HCMV was recovered by electroporation of BAC DNA into MRC-5 fibroblasts as described by Wille et al. (22). Cell-free HCMV stocks were produced by infecting HFF or HFFFtet at 2 plaque-forming unit (PFU) per cell. At 8-10 days post infection (when cells were still visually intact), culture supernatants were harvested, and cellular contaminants were removed by centrifugation at 1,000 × g for 10min, and again at 6,000 × g for 10min. Stocks were judged cell-free by the lack of calnexin, and actin in western blot analyses, and then stored at -80°C. Freeze/thaw cycles were avoided. Plaque-forming-units were determined by plating a series of 10-fold dilutions of each stock on replicate cultures of HFF for 2h at 37° C, and replacing the inoculum with a DMEM supplemented with 5% FBS, and 0.6% SeaPlaque agarose (to limit cell free spread). Plaques were counted by light microscopy 3 weeks after infection.

### Heterologous UL74(gO) recombinant HCMV

A two step BAC recombineering process was performed as previously described (33). In the first step, endogenous UL74 ORF from start codon to stop codon of both TR and ME was replaced by a selectable marker. Briefly, overnight cultures of SW102 *E. coli* containing either the BAC clone of TR or ME were grown at 32° C until OD600=0.55. Recombination genes were induced by incubating at 42° C for 15 mins. Purified PCR product containing the selectable marker cassette, KanR/LacZ/RpsL flanked by sequences homologous to 80 base pairs upstream and downstream of the TR or ME UL74 ORF, was electroporated into the bacteria, cultures were recovered for 1h at 32° C, and then selected on media containing kanamycin (15ug/ml), IPTG (50uM), X-gal (20ug/ml) and chloramphenicol (12.5ug/ml). First-step primer sequences were; TR: 5′-

CTTGGTGGACTATGCTTAACGCTCTCATTCTCATGGGAGCTTTTTGTATCGTATTACGACATTGCTGTTTCCAGAACTCCTGTGACGGAAGATCACTTCG-3′; 5′-

CGACCAGAATCAGCAGTGAGTACACGCAGGCAAGCCAAACCACAAGGCAGACGGACGGTGCGGGGTCTCCTCCTCTGTCCTGAGGTTCTTATGGCTCTTG-3′; ME: 5′-

CCTGGTGGACTATGCTTAACGCTCTCATTCTGATGGGAGCTTTTTGTATCGTATTACGACATTGCTGCTTCCAGAACTCCTGTGACGGAAGATCACTTCG-3′; 5′-

CGACCAGAATCAGCAGTGAGTACACGCAGGCAAACCAAACCACAAGGCAGACGGACGGTGCGGGGTCTCCTCCTCTGTACTGAGGTTCTTATGGCTCTTG-3′

In the second step, the selectable marker cassette in the TR and ME first-step intermediate BACs was replaced with the UL74(gO) sequence from the heterologous strain. Briefly, *E. coli* were prepared for recombination as described for step one above, and electroporated with purified PCR products containing the UL74 ORF from TR or ME strain flanked by sequence homologous to 80 base pairs upstream and downstream of the opposite strain. Transformed *E. coli* were selected for the removal of the KanR/LacZ/RpsL cassette by growth on media containing streptomycin (1.5mg/ml), IPTG (50uM), X-gal (20ug/ml) and chloramphenicol (12.5ug/ml). Primers used to generate the second-step PCR produce were, TR UL74 into ME: 5′-GCCTGGTGGACTATGCTTAACGCTCTCATTCTGATGGGAGCTTTTTGTATCGTATTACGACA TTGCTGCTTCCAGAACTTTACTGCAACCACCACCAAAG-3′, and 5′-

CGACCAGAATCAGCAGTGAGTACACGCAGGCAAACCAAACCACAAGGCAGACGGACGGTGCGGGGTCTCCTCCTCTGTAATGGGGAGAAAAGGAGAGATG-3′. ME UL74 into TR: 5′-

GGCTTGGTGGACTATGCTTAACGCTCTCATTCTCATGGGAGCTTTTTGTATCGTATTACGACATTGCTGTTTCCAGAACTTTACTGCGACCACCACCAAA-3′, and 5′-

CAGAATCAGCAGTGAGTACACGCAGGCAAGCCAAACCACAAGGCAGACGGACGGTGCGGGGTCTCCTCCTCTGTCATGGGGAAAAAAGAGATGATAATGG

The final heterologous UL74(gO) recombinants were verified by Sanger sequencing PCR products using the following primers; TRMEgO: 5′-

GATGATTTTTACAAGGCACATTGTACATC-3′, and 5′-AACTAGGTCGTCTTGGAAGC-3′, METRgO: 5′-CTCACAATGATTTTTACAATGCG-3′, and 5′-AACTAGGTCGTCTTGGAAGC-3′.

### Antibodies

Rabbit polyclonal anti-peptide antibodies specific for TBgO and MEgO were described previously (30). Rabbit polyclonal antibodies specific for UL148 were described (32). Rabbit polyclonal, anti-peptide antibodies against gH, gL, UL130 and UL131 were provided by David Johnson (Oregon Health and Sciences University, Portland, OR, USA) (45). Anti-UL128 monoclonal antibodies (mAb) 4B10 were provided by Tom Shenk (Princeton University, Princeton, NJ, USA) (46). mAbs directly against major capsid protein (MCP) 28-4 and gB 27-156 were provided by Bill Britt (47, 48). mAb (CH160) against CMV immediate early protein1 and immediate early protein2 (IE1/IE2) was purchased from Abcam (Cambridge, MA, USA).

### Immunoblot

HCMV-infected cells or cell free virions were solubilized in 2% SDS/ 20mM Tris-buffered saline (TBS) (pH 6.8). Insoluble material was cleared by centrifugation at 16,000 × g for 15 min and then extracts were boiled for 10 min. For endoglycosidase H (endo H) or peptide N-glycosidase F (PNGase F) treatment assay, proteins were extracted in 1% Triton X-100 (TX100), 0.5% sodium deoxycholate (DOC) in 20 mM Tris (pH 6.8), 100 mM NaCl (TBS-TX/DOC). Extracts were clarified by centrifugation 16,000 × g for 15 min and treated with with endo H or PNGase F according to manufacturer′s instructions (New England BioLabs). For reducing blots, dithiothereitol (DTT) were added to extracts to a final concentration of 25 mM. After separation by SDS-PAGE, proteins were transferred to polyvinylidene difluoride (PVDF) membranes (Whatman) in a buffer containing 10 mM NaHCO_3_, 3 mM Na_2_CO_3_ (pH 9.9) plus 10% methanol. Transferred proteins were probed with mAbs or rabbit polyclonal antibodies, anti-rabbit or anti-mouse secondary antibodies conjugated with horseradish peroxidase (Sigma-Aldrich), and Pierce ECL-Western Blotting Substrate (ThermoFisher Scientific). Chemiluminescence was detected using a Bio-Rad ChemiDoc MP imaging system.

### Radiolabeling proteins

Cell cultures were incubated in labeling medium (met/cys-free DMEM + 2% dialyzed FBS) lacking methionine and cysteine) for 2 h at 37 °C, then [^35^S] methionine-cysteine was added to 1 mCi/ml (EasyTag Express 35S Protein Labeling Mix; Perkin Elmer). For chase experiments, label medium was removed and cultures were washed twice in DMEM + 2% FBS supplemented with a 10-fold excess of nonradioactive methionine and cysteine, then incubated in this medium for the indicated time.

### Immunoprecipitation

Cell extracts were harvested in TBS-TX/DOC supplemented with 0.5% bovine serum albumin (BSA) and 1mM phenylmethylsulfonyl fluoride (PMSF), clarified by centrifugation at 16,000 × g for 15 min, adjusted to 2% SDS, 30 mM DTT and heated at 75 °C for 15 min. The extracts were then diluted 35-fold with TBS-TX/DOC supplemented with 0.5% BSA, and 10 mM iodoacetamide, incubated on ice for 15 min and pre-cleared with protein A-agarose beads (Invitrogen/ThermoFisher Scientific) for at 4 °C for 2 h. Immunoprecipition reactions were set up with specific antibodies and protein A-agarose beads and incubated overnight at 4°C. Protein A-agarose beads were washed 3 times with TBS-TX/DOC and proteins were eluted with 2% SDS, 30 mM DTT in TBS at RT for 15 min, followed by 75 °C for 10 min. Eluted proteins were separated by SDS-PAGE and analyzed with a Typhoon FLA-9500 imager (GE Healthcare Life Sciences). Band densities were determined using ImageJ version 1.48 software.

### Quantitative PCR

Viral genomes were determined as described (25). Briefly, Cell-free HCMV stocks were treated with DNase I before extraction of viral genomic DNA (PureLink Viral RNA/DNA minikit; Life Techonologies/ThermoFisher Scientific). Primers specific for sequences within UL83 were used with the MyiQ real-time PCR detection system (Bio-Rad).

### Superinfection of HCMV-infected cells with replication-defective adenovirus vectors

Construction of Ad vectors expressing MEgO or GFP was described (30). At 2 days post HCMV infection, cells were superinfected with 20 PFU/cell of AdMEgO or AdGFP. 6 days later, cell-free HCMV was collected from the supernatant culture by centrifugation, and cells were harvested for immunoblot.

## ACKNOWLEDGMENTS

We are grateful to Bill Britt, David Johnson, Jay Nelson, and Tom Shenk, for generously supplying HCMV BAC clones, antibodies, and cell lines as indicated in the Material and Methods, and members of the Ryckman laboratory for support, and insightful discussions.

This work was supported by grant from the National Institutes of Health to B.J.R (R01AI097274).

Experiments were designed by B.J.R., LZ, and M.Z., and performed by L.Z., and M.Z. Data were analyzed, and manuscript was prepared by B.J.R., L.Z., J.P.K. and, R.J.S.

## REFERENCES

1. Boppana SB, Fowler KB, Pass RF, Rivera LB, Bradford RD, Lakeman FD, Britt WJ. 2005. Congenital cytomegalovirus infection: association between virus burden in infancy and hearing loss. J Pediatr 146:817–823.

2. Britt WJ. 2008. Manifestations of human cytomegalovirus infection: proposed mechanisms of acute and chronic disease., p. In T.E. S, M.F. S (ed), Human Cytomegalovirus. Current Topics in Microbiology and Immunology 325,

3. Griffiths P, Baraniak I, Reeves M. 2015. The pathogenesis of human cytomegalovirus. J Pathol 235:288–297.

4. Streblow DN, Orloff SL, Nelson JA. 2007. Acceleration of allograft failure by cytomegalovirus. Curr Opin Immunol 19:577–582.

5. Cannon MJ, Hyde TB, Schmid DS. 2011. Review of cytomegalovirus shedding in bodily fluids and relevance to congenital cytomegalovirus infection. Rev Med Virol 21:240–255.

6. Plachter B, Sinzger C, Jahn G. 1996. Cell types involved in replication and distribution of human cytomegalovirus. Adv Virus Res 46:195–261.

7. Sinzger C, Grefte A, Plachter B, Gouw AS, The TH, Jahn G. 1995. Fibroblasts, epithelial cells, endothelial cells and smooth muscle cells are major targets of human cytomegalovirus infection in lung and gastrointestinal tissues. J Gen Virol 76:741–750.

8. Sinzger C, Jahn G. 1996. Human cytomegalovirus cell tropism and pathogenesis. Intervirology 39:302–319.

9. Sinzger C, Digel M, Jahn G. 2008. Cytomegalovirus cell tropism. Curr Top Microbiol Immunol 325:63–83.

10. Xu S, Schafer X, Munger J. 2016. Expression of Oncogenic Alleles Induces Multiple Blocks to Human Cytomegalovirus Infection. J Virol 90:4346–4356.

11. Smith JD. 1986. Human cytomegalovirus: demonstration of permissive epithelial cells and nonpermissive fibroblastic cells in a survey of human cell lines. J Virol 60:583–588.

12. Connolly SA, Jackson JO, Jardetzky TS, Longnecker R. 2011. Fusing structure and function: a structural view of the herpesvirus entry machinery. Nat Rev Microbiol 9:369–381.

13. Campadelli-Fiume G, Collins-McMillen D, Gianni T, Yurochko AD. 2016. Integrins as Herpesvirus Receptors and Mediators of the Host Signalosome. Annu Rev Virol 3:215–236.

14. Heldwein EE. 2016. gH/gL supercomplexes at early stages of herpesvirus entry. Curr Opin Virol 18:1–8.

15. Ciferri C, Chandramouli S, Donnarumma D, Nikitin PA, Cianfrocco MA, Gerrein R, Feire AL, Barnett SW, Lilja AE, Rappuoli R, Norais N, Settembre EC, Carfi A. 2015. Structural and biochemical studies of HCMV gH/gL/gO and Pentamer reveal mutually exclusive cell entry complexes. Proc Natl Acad Sci U S A 112:1767–1772.

16. Stegmann, Abdellatif, Sampaio L, Walther., Sinzger. 2017. Importance of Highly Conserved Peptide Sites of Human Cytomegalovirus gO for Formation of the gH/gL/gO Complex. J Virol 91

17. Hahn G, Revello MG, Patrone M, Percivalle E, Campanini G, Sarasini A, Wagner M, Gallina A, Milanesi G, Koszinowski U, Baldanti F, Gerna G. 2004. Human cytomegalovirus UL131-128 genes are indispensable for virus growth in endothelial cells and virus transfer to leukocytes. J Virol 78:10023–10033.

18. Ryckman BJ, Jarvis MA, Drummond DD, Nelson JA, Johnson DC. 2006. Human cytomegalovirus entry into epithelial and endothelial cells depends on genes UL128 to UL150 and occurs by endocytosis and low-pH fusion. J Virol 80:710–722.

19. Wang D, Shenk T. 2005. Human cytomegalovirus UL131 open reading frame is required for epithelial cell tropism. J Virol 79:10330–10338.

20. Luo MH, Schwartz PH, Fortunato EA. 2008. Neonatal neural progenitor cells and their neuronal and glial cell derivatives are fully permissive for human cytomegalovirus infection. J Virol 82:9994–10007.

21. Gerna G, Percivalle E, Lilleri D, Lozza L, Fornara C, Hahn G, Baldanti F, Revello MG. 2005. Dendritic-cell infection by human cytomegalovirus is restricted to strains carrying functional UL131-128 genes and mediates efficient viral antigen presentation to CD8+ T cells. J Gen Virol 86:275–284.

22. Wille PT, Knoche AJ, Nelson JA, Jarvis MA, Johnson DC. 2010. A human cytomegalovirus gO-null mutant fails to incorporate gH/gL into the virion envelope and is unable to enter fibroblasts and epithelial and endothelial cells. J Virol 84:2585–2596.

23. Jiang XJ, Adler B, Sampaio KL, Digel M, Jahn G, Ettischer N, Stierhof YD, Scrivano L, Koszinowski U, Mach M, Sinzger C. 2008. UL74 of human cytomegalovirus contributes to virus release by promoting secondary envelopment of virions. J Virol 82:2802–2812.

24. Laib Sampaio K, Stegmann C, Brizic I, Adler B, Stanton RJ, Sinzger C. 2016. The contribution of pUL74 to growth of human cytomegalovirus is masked in the presence of RL13 and UL128 expression. J Gen Virol 97:1917–1927.

25. Zhou M, Lanchy JM, Ryckman BJ. 2015. Human cytomegalovirus gH/gL/gO promotes the fusion step of entry into all cell types whereas gH/gL/UL128-131 broadens virus tropism through a distinct mechanism. J Virol

26. Kabanova A, Marcandalli J, Zhou T, Bianchi S, Baxa U, Tsybovsky Y, Lilleri D, Silacci-Fregni C, Foglierini M, Fernandez-Rodriguez BM, Druz A, Zhang B, Geiger R, Pagani M, Sallusto F, Kwong PD, Corti D, Lanzavecchia A, Perez L. 2016. Platelet-derived growth factor-alpha receptor is the cellular receptor for human cytomegalovirus gHgLgO trimer. Nat Microbiol 2016

27. Stegmann C, Hochdorfer D, Lieber D, Subramanian N, Stöhr D, Laib Sampaio K, Sinzger C. 2017. A derivative of platelet-derived growth factor receptor alpha binds to the trimer of human cytomegalovirus and inhibits entry into fibroblasts and endothelial cells. PLoS Pathog 13:e1006273.

28. Wu Y, Prager A, Boos S, Resch M, Brizic I, Mach M, Wildner S, Scrivano L, Adler B. 2017. Human cytomegalovirus glycoprotein complex gH/gL/gO uses PDGFR-α as a key for entry. PLoS Pathog 13:e1006281.

29. Nogalski MT, Chan GC, Stevenson.EV, Collins-McMillen DK, Yurochko AD. 2013. The HCMV gH/gL/UL128-131 Complex Triggers the Specific Cellular Activation Required for Efficient Viral Internalization into Target Monocytes. PLoS Pathog 9:e1003463.

30. Zhou M, Yu Q, Wechsler A, Ryckman BJ. 2013. Comparative analysis of gO isoforms reveals that strains of human cytomegalovirus differ in the ratio of gH/gL/gO and gH/gL/UL128-131 in the virion envelope. J Virol 87:9680–9690.

31. Rasmussen L, Geissler A, Cowan C, Chase A, Winters M. 2002. The genes encoding the gCIII complex of human cytomegalovirus exist in highly diverse combinations in clinical isolates. J Virol 76:10841–10848.

32. Li G, Nguyen CC, Ryckman BJ, Britt WJ, Kamil JP. 2015. A viral regulator of glycoprotein complexes contributes to human cytomegalovirus cell tropism. Proc Natl Acad Sci U S A 112:4471–4476.

33. Stanton RJ, Baluchova K, Dargan DJ, Cunningham C, Sheehy O, Seirafian S, McSharry BP, Neale ML, Davies JA, Tomasec P, Davison AJ, Wilkinson GW. 2010. Reconstruction of the complete human cytomegalovirus genome in a BAC reveals RL13 to be a potent inhibitor of replication. J Clin Invest 120:3191–3208.

34. Wiertz EJ, Jones TR, Sun L, Bogyo M, Geuze HJ, Ploegh HL. 1996. The human cytomegalovirus US11 gene product dislocates MHC class I heavy chains from the endoplasmic reticulum to the cytosol. Cell 84:769–779.

35. Renzette N, Bhattacharjee B, Jensen JD, Gibson L, Kowalik TF. 2011. Extensive genome-wide variability of human cytomegalovirus in congenitally infected infants. PLoS Pathog 7:e1001344.

36. Renzette N, Gibson L, Bhattacharjee B, Fisher D, Schleiss MR, Jensen JD, Kowalik TF. 2013. Rapid Intrahost Evolution of Human Cytomegalovirus Is Shaped by Demography and Positive Selection. PLoS Genet 9:e1003735.

37. Sijmons S, Thys K, Mbong Ngwese M, Van Damme E, Dvorak J, Van Loock M, Li G, Tachezy R, Busson L, Aerssens J, Van Ranst M, Maes P. 2015. High-throughput analysis of human cytomegalovirus genome diversity highlights the widespread occurrence of gene-disrupting mutations and pervasive recombination. J Virol

38. Dargan DJ, Douglas E, Cunningham C, Jamieson F, Stanton RJ, Baluchova K, McSharry BP, Tomasec P, Emery VC, Percivalle E, Sarasini A, Gerna G, Wilkinson GW, Davison AJ. 2010. Sequential mutations associated with adaptation of human cytomegalovirus to growth in cell culture. J Gen Virol 91:1535–1546.

39. Murrell I, Wilkie GS, Davison AJ, Statkute E, Fielding CA, Tomasec P, Wilkinson GW, Stanton RJ. 2016. Genetic Stability of Bacterial Artificial Chromosome-Derived Human Cytomegalovirus during Culture In Vitro. J Virol 90:3929–3943.

40. Murrell I, Tomasec P, Wilkie GS, Dargan DJ, Davison AJ, Stanton RJ. 2013. Impact of sequence variation in the UL128 locus on production of human cytomegalovirus in fibroblast and epithelial cells. J Virol 87:10489–10500.

41. Balázs Z, Tombácz D, Szűcs A, Csabai Z, Megyeri K, Petrov AN, Snyder M, Boldogkői Z. 2017. Long-Read Sequencing of Human Cytomegalovirus Transcriptome Reveals RNA Isoforms Carrying Distinct Coding Potentials. Sci Rep 7:15989.

42. Gatherer D, Seirafian S, Cunningham C, Holton M, Dargan DJ, Baluchova K, Hector RD, Galbraith J, Herzyk P, Wilkinson GW, Davison AJ. 2011. High-resolution human cytomegalovirus transcriptome. Proc Natl Acad Sci U S A 108:19755–19760.

43. Kalser J, Adler B, Mach M, Kropff B, Puchhammer-Stöckl E, Görzer I. 2017. Differences in Growth Properties among Two Human Cytomegalovirus Glycoprotein O Genotypes. Front Microbiol 8:1609.

44. Murphy E, Yu D, Grimwood J, Schmutz J, Dickson M, Jarvis MA, Hahn G, Nelson JA, Myers RM, Shenk TE. 2003. Coding potential of laboratory and clinical strains of human cytomegalovirus. Proc Natl Acad Sci U S A 100:14976–14981.

45. Ryckman BJ, Rainish BL, Chase MC, Borton JA, Nelson JA, Jarvis MA, Johnson DC. 2008. Characterization of the human cytomegalovirus gH/gL/UL128-131 complex that mediates entry into epithelial and endothelial cells. J Virol 82:60–70.

46. Wang D, Shenk T. 2005. Human cytomegalovirus virion protein complex required for epithelial and endothelial cell tropism. Proc Natl Acad Sci U S A 102:18153–18158.

47. Schoppel K, Hassfurther E, Britt W, Ohlin M, Borrebaeck CA, Mach M. 1996. Antibodies specific for the antigenic domain 1 of glycoprotein B (gpUL55) of human cytomegalovirus bind to different substructures. Virology 216:133–145.

48. Chee M, Rudolph SA, Plachter B, Barrell B, Jahn G. 1989. Identification of the major capsid protein gene of human cytomegalovirus. J Virol 63:1345–1353.

